# Tissue-resident memory CD8 T-cell responses elicited by a single injection of a multi-target COVID-19 vaccine

**DOI:** 10.1101/2020.08.14.240093

**Authors:** V. Gauttier, A. Morello, I. Girault, C. Mary, L. Belarif, A. Desselle, E. Wilhelm, T. Bourquard, S. Pengam, G. Teppaz, V. Thepenier, K. Biteau, E. De Barbeyrac, D. Kiepferlé, B. Vasseur, FX. Le Flem, D. Debieuvre, D. Costantini, N. Poirier

**Author notes:** These authors contributed equally to the first authorship of this study (V. Gauttier, A. Morello, I. Girault, C. Mary). Correspondence should be addressed to: Dr. Nicolas Poirier, OSE Immunotherapeutics, 22 Boulevard Benoni Goullin, 44200 Nantes, France.

## Abstract

The COVID-19 pandemic is caused by severe acute respiratory syndrome coronavirus-2 (SARS-CoV-2) which enters the body principally through the nasal and larynx mucosa and progress to the lungs through the respiratory tract. SARS-CoV-2 replicates efficiently in respiratory epithelial cells motivating the development of alternative and rapidly scalable vaccine inducing mucosal protective and long-lasting immunity. We have previously developed an immunologically optimized multi-neoepitopes-based peptide vaccine platform which has already demonstrated tolerance and efficacy in hundreds of lung cancer patients. Here, we present a multi-target CD8 T cell peptide COVID-19 vaccine design targeting several structural (S, M, N) and non-structural (NSPs) SARS-CoV-2 proteins with selected epitopes in conserved regions of the SARS-CoV-2 genome. We observed that a single subcutaneous injection of a serie of epitopes induces a robust immunogenicity in-vivo as measured by IFNγ ELIspot. Upon tetramer characterization we found that this serie of epitopes induces a strong proportion of virus-specific CD8 T cells expressing CD103, CD44, CXCR3 and CD49a, the specific phenotype of tissue-resident memory T lymphocytes (Trm). Finally, we observed broad cellular responses, as characterized by IFNγ production, upon restimulation with structural and non-structural protein-derived epitopes using blood T cells isolated from convalescent asymptomatic, moderate and severe COVID-19 patients. These data provide insights for further development of a second generation of COVID-19 vaccine focused on inducing lasting Th1-biased memory CD8 T cell sentinels protection using immunodominant epitopes naturally observed after SARS-CoV-2 infection resolution.

**Statement of Significance:** Humoral and cellular adaptive immunity are different and complementary immune defenses engaged by the body to clear viral infection. While neutralizing antibodies have the capacity to block virus binding to its entry receptor expressed on human cells, memory T lymphocytes have the capacity to eliminate infected cells and are required for viral clearance. However, viruses evolve quickly, and their antigens are prone to mutations to avoid recognition by the antibodies (phenomenon named ‘antigenic drift’). This limitation of the antibody-mediated immunity could be addressed by the T-cell mediated immunity, which is able to recognize conserved viral peptides from any viral proteins presented by virus-infected cells. Thus, by targeting several proteins and conserved regions on the genome of a virus, T-cell epitope-based vaccines are less subjected to mutations and may work effectively on different strains of the virus. We designed a multi-target T cell-based vaccine containing epitope regions optimized for CD8+ T cell stimulation that would drive long-lasting cellular immunity with high specificity, avoiding undesired effects such as antibody-dependent enhancement (ADE) and antibody-induced macrophages hyperinflammation that could be observed in subjects with severe COVID-19. Our in-vivo results showed that a single injection of selected CD8 T cell epitopes induces memory viral-specific T-cell responses with a phenotype of tissue-resident memory T cells (Trm). Trm has attracted a growing interest for developing vaccination strategies since they act as immune sentinels in barrier tissue such as the respiratory tract and the lung. Because of their localization in tissues, they are able to immediately recognize infected cells and, because of their memory phenotypes, they rapidly respond to viral infection by orchestrating local protective immune responses to eliminate pathogens. Lastly, such multiepitope-based vaccination platform uses robust and well-validated synthetic peptide production technologies that can be rapidly manufactured in a distributed manner.

## Introduction

COVID-19, the infectious disease caused by the zoonotic coronavirus SARS-CoV-2, is a global pandemic which has infected more than 16 million of individuals and already caused up to 650 000 deaths worldwide as of end of July 2020. The severity of the COVID-19 outbreak highlights the need to develop therapeutic interventions including fast and efficacious prophylactic vaccines. Previous severe outbreaks of coronavirus (CoVs) have emerged including Severe acute respiratory syndrome CoV (SARS-CoV-1) in 2002 and Middle East Respiratory Syndrome CoV (MERS-CoV) in 2012. Learnings from current (SARS-CoV-2) and previous (SARS-CoV-1, MERS-CoV) infections argue that the body develops humoral and cellular immune responses against this type of respiratory viruses but that antibody responses are short-term^1–5^ in contrast to the cellular immunity which is still observed 11 and 17 years after the infection^6,7^.

Current and previous CoVs vaccine strategies have been almost exclusively focused on eliciting a humoral immune response, particularly anti-Spike neutralizing IgG antibody. However, the generation of non-neutralizing antibody responses^8^, insufficient antibody titers^9^, Th2-biased immune response or glycosylation changes in the IgG Fc tail^10^ may be associated with vaccine failure, and in the worst case scenario may enhance disease upon viral exposure, either through the induction of enhanced pulmonary macrophage-mediated hyper-inflammation^11^, or Fc receptor-mediated antibody-dependent enhancement (ADE)^12^. Th1-biasing immunization using CD8 T cells optimized peptide vaccination may offer an important alternative and complementary approach with a history of safe administration, may be developed and updated rapidly, and should avoid safety pitfalls in the pursuit of a COVID-19 vaccine^8,13,14^.

The discovery of memory T lymphocytes resident in diverse tissues, in particular mucosal and barrier tissues, has highlighted the importance of site-specific responses and continuous surveillance mediated by a specific tissue-resident memory T cell population (Trm)^15–17^. A couple of studies demonstrated that after induction Trm migrate and reside in the lung, skin, and gut long after infection resolution and provide localized protective immunity and immunosurveillance in tissues^18–21^. Trm represents an attractive target population and a growing interest for developing vaccine strategies since they act as immune sentinels in mucosal and barrier tissue and rapidly respond to infection by orchestrating local protective immune responses to eliminate pathogens^22^. Previous CD8 T cell-based vaccines against SARS-CoV-1^23^ and influenza A^24^ viruses showed lasting virus-specific memory CD8 T cells induction in the spleen, lung and bronchoalveolar fluids (BALs) and protection of mice from lethal SARS-CoV-1 or influenza challenges. Expression of chemokine receptors, such as CXCR3, is critical for CD8+ Trm to populate the airways after vaccination and protection against influenza A viruses^25^. More recently, a Trm-inducing HIV vaccine durably prevented mucosal infection in non-human primates even with lower neutralizing antibody titers^26^. Altogether, these data emphasize the interest of tissue-resident memory viral-specific CD8 T cell generation upon vaccination for optimal protection against airways viruses.

Stimulation of a proper immune response that leads to protection is highly dependent on presentation of epitopes to circulating T-cells via the HLA complex. MEMOPI^®^ is a robust vaccine platform based on selection and immunogenicity optimization of HLA-restricted peptides (neo-epitope) technologies^27,28^ and formulation^29^ for multi-epitopes and targets combination with a pan-DR helper epitope (PADRE) providing help for memory CD8 T cell generation^30^. A combination of multi-target antitumor neo-epitopes, Tedopi^®^, based on this platform already demonstrated good safety profile and efficacy in clinical Phase 2 trial^31^ and more recently successfully validated in the first step of a Phase 3 clinical testing in lung cancer paients^32^. While induction of mucosal immunity using parenteral administration of conventional virus vaccine technologies was challenging, we observed that subcutaneous injection of our neo-epitopes multi-target cancer vaccine promotes Th1-biased antigen-specific memory CD8 T cell responses in the lung and BALs of vaccinated mice in the absence of tumor. Based on this original preclinical observation and the significant survival increase measured during clinical trials in lung cancer patients correlating with epitope responses, we generated and screened individually a large number of immuno-dominant SARS-CoV-2 epitopes, and their neo-epitopes generated using artificial intelligence (AI) algorithms, covering all sequenced circulating SARS-CoV-2 strains and derived from 11 structural (including Spike) and non-structural proteins with significant homology with previous SARS-CoV-1 virus.

## Results

### CD8 T-cell epitopes selection and neo-epitopes design

Previous research in SARS-CoV-1 suggests that the structural Spike (S) protein is one of the main antigenic component responsible for inducing the host immune responses^33^. However recent evaluation of asymptomatic, moderate and severe convalescent COVID-19 patients showed broad and robust T-cell responses not only on Spike but also on membrane (M), nucleocapsid (N) proteins and several ORFs non-structural proteins (nsp)^34–37^. Deep sequencing data of tens of thousands SARS-CoV-2 genomes from all over the world identified regions that have remained largely invariant to date, and others that have already accumulated significant diversity with several hundreds of point mutations (SNPs) in some key viral proteins, such as the Spike glycoprotein which also displayed a large number of recurrent mutations and several homoplasic sites highlighting a possible convergent evolution and adaptation of SARS-CoV-2 to the human host^38–41^. COVID-19 vaccines reliance on a single antigen (Spike) bearing recurrent mutations and homoplasic sites, as occurring with Spike monovalent vaccines under development, is not without risk of antigen drift, selection pressure and immune evasion^42^.

Using bioinformatics approaches of immune deconvolution to identify T cell epitopes intersected with predicted SARS-CoV-2 T-cell epitopes reported in early manuscripts^43–49^, population HLA diversity, protein sequence similarity and immunogenicity observed with previous CoVs homologues and the number of copies per virion of SARS-COV-1^50,51^ (estimated to approximatively 100 copies for S, 1000 for M, 2000 for N and only 20 for E proteins), we identified and selected 55 HLA-A*0201-restricted T-cell epitopes (9-10 mer peptides) derived from 11 of the 29 SARS-CoV-2 proteins in human cells^52^: 3 out of the 4 structural proteins (S, M, N), the largest accessory proteins (ORF3a) and 7 out of the 16 ORF1a/b non-structural proteins (nsp3, nsp4, nsp6, nsp12, nsp13, nsp14, nsp16) (**Figure 1** and **Table 1**). Based on our knowledge of key fixed-anchor positions to enhance HLA binding and increase their immunogenicity potential, we designed 400 mutated sequences for each individual peptide resulting in 22 000 total analyzed sequences of the 55 selected epitopes. We first screened these potential peptides using *in-silico* bioinformatic analyses (*e.g.* IEDB Immune epitope database, netMHCpan EL 4.0 algorithm) and a first series of the most optimized mutant for each epitope was selected (neo-epitopes A). In parallel, 22000 *in silico* HLA-A*0201-peptide docking models were generated using computational tools and analyzed using newly developed proprietary artificial intelligence (AI) algorithms based on binding energy calculations and reduction (MabSilico, France). A second series of the most optimized mutant for each epitope was selected (neo-epitopes B). Finally, SARS-CoV-2 genetic evolution analyses through the alignment of 46723 sequences (*https://macman123.shinyapps.io/ugi-scov2-alignment-screen/*) identified recurrent mutation (SNPs) and homoplasic site in SARS-CoV-2 genomes isolated globally, particularly in the Spike protein which contain the D614G mutation and which identified the new dominant SARS-CoV-2 variant emerged in February in Europe, then spread worldwide and became the most prevalent form^62,63^. We eliminated T cell epitopes with recurrent mutation and homoplasic site in order to cover all circulating SARS-CoV-2 strains and anticipate future evolution of the virus in hotspot mutation regions.

**Figure 1:**
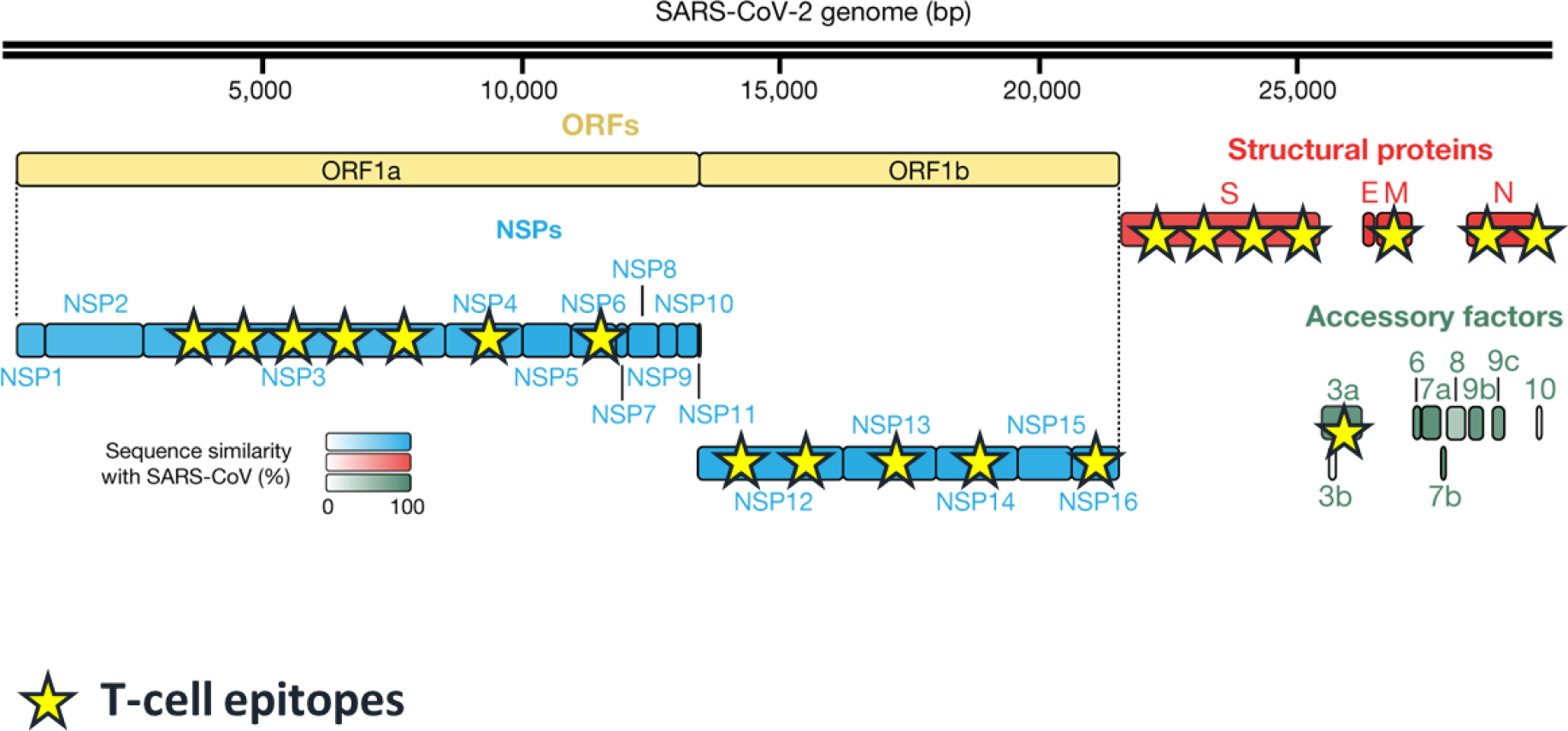
T-cell epitopes location in SARS-CoV-2 genome. SARS-CoV-2 genome annotation by the Krogran lab^52^ and schematic representation of T-cell epitopes location in each encoded proteins. n=4 structural proteins; n=16 non-structural proteins (NSPs); n=9 accessory factors.

**Table 1:**
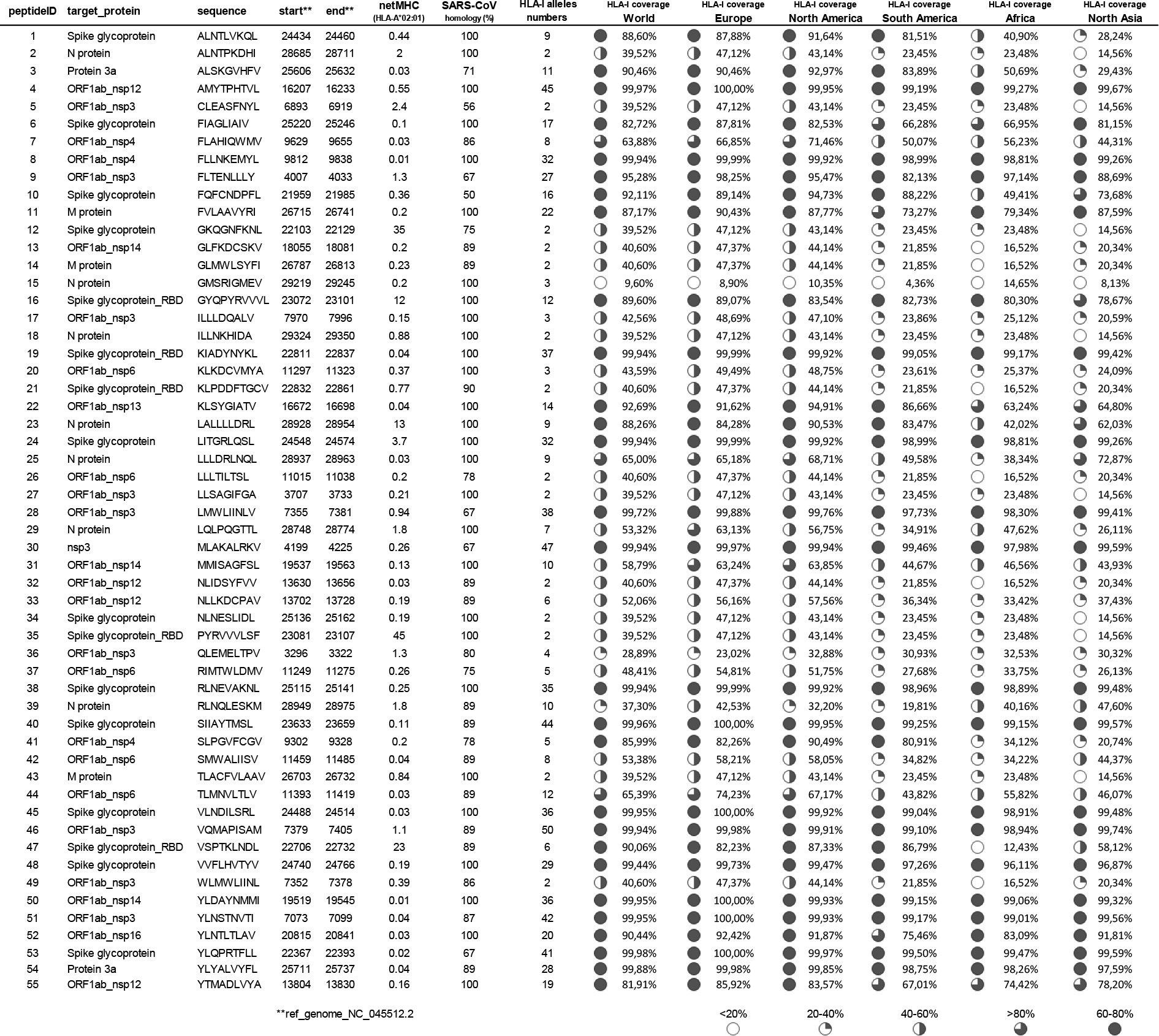
T-cell WT epitopes. HLA-I alleles numbers and regional HLA-I coverage were determined using IEDB public database and netMHCpan score < 1.

### CD8 T-cell epitopes elicit Tissue-resident memory (Trm) viral-specific T cells in-vivo

134 WT and mutated peptides (neoepitopes A and B) were produced using synthetic peptide synthesis (Proteogenix, France). HLA-A2 binding property characterization at 37°C, using UV peptide exchange assay on HLA-A*0201 monomer, showed that the majority of selected WT epitopes binds to HLA-A2 with good efficacy (**Figure 2A**) as compared to our MEMOPI^®^ internal positive neoepitope control (mutated peptide with increased HLA-A*0201 binding and *in-vivo* immunogenicity). HLA-A2 binding was increased with several neoepitopes A and/or B, particularly when the corresponding WT peptide showed weak (< 15%: peptides 2, 9, 12, 24, 29 and 35) or intermediate (15-30%: peptides 4, 14, 16, 18 and 48) HLA-A2 stability at 37°C. Similarly, several neoepitopes A increased peptide binding to HLA-A2 expressing human cells (TAP-deficient human cell line (T2) lacking the ability to transport peptide fragments to the endoplasmic reticulum to form stable pMHCI) particularly when the corresponding WT peptide showed low (< 30%: peptides 34, 39) or intermediate (30-60%: peptides 4, 5, 21, 26, 38) HLA-A2 stability at 37°C (**Figure 2B**). Altogether, bioinformatic and AI-based mutation increased peptides stability with HLA-A2 (recombinant or expressed by cells) for 52% (26/50) of evaluated CD8 T-cell epitopes.

**Figure 2:**
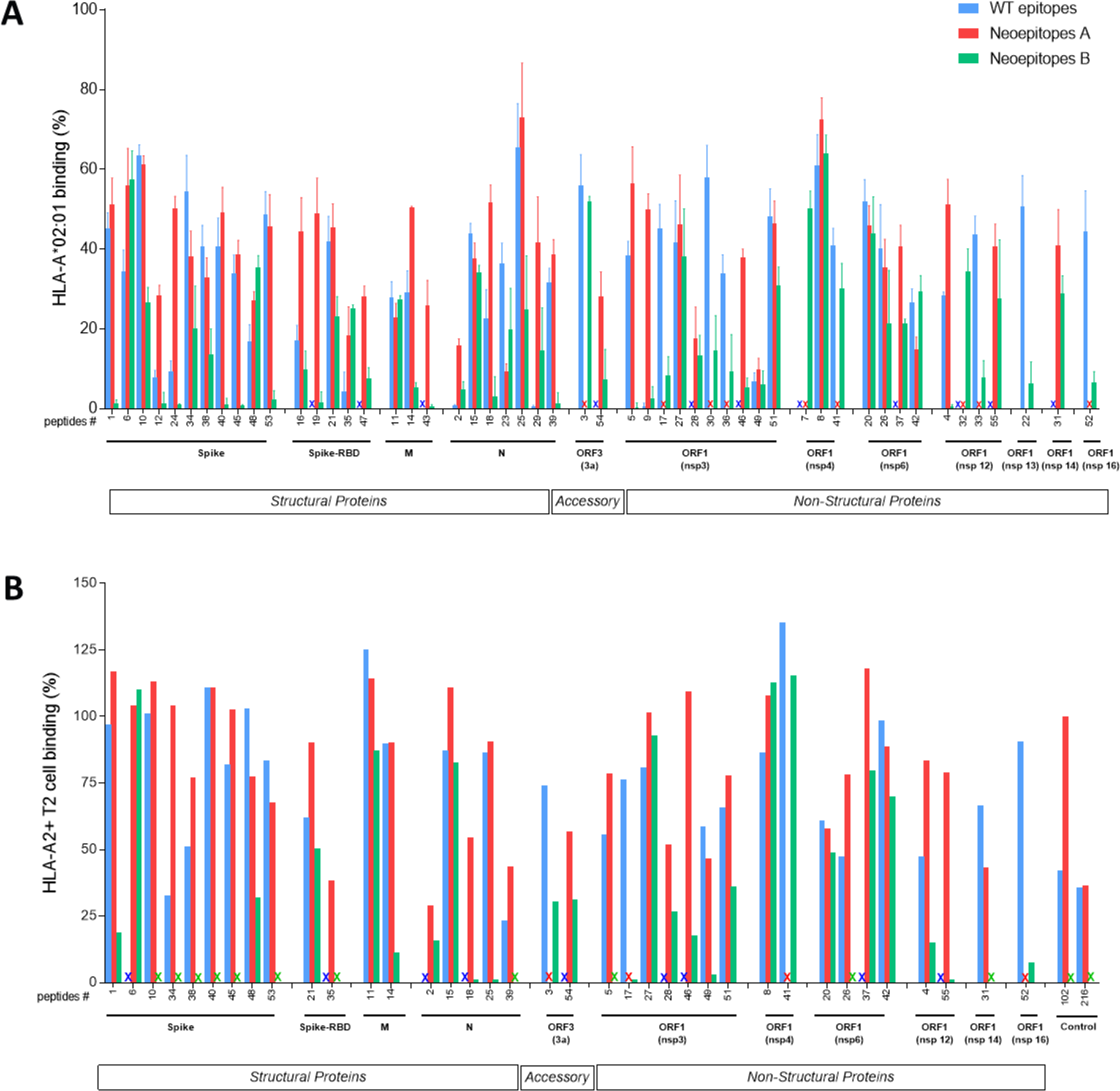
HLA-A2 binding characterization of WT and mutated T-cell epitopes. (**A**) WT and mutated peptides were incubated with HLA-A*02:01 monomer, exposed to UV for peptide exchange and then HLA-peptide complexes stability at 37°C was measured by ELISA. Data are mean +/- SEM (n=4) expressed as percentage of binding relative to an internal MEMOPI^®^ positive control neoepitope. (**B**) WT and mutated peptides (25μM) binding to TAP-deficient human cell line (T2) expressing HLA-A2. Data are expressed as percentage of binding relative to an internal MEMOPI^®^ positive control neoepitope. X mark indicates that the peptide was not tested in the assay.

Finally, 60 peptides (at least one WT or mutated peptides for each T-cell epitopes outside homoplasic site) has been selected based on HLA-A2 binding, peptide stability and SARS-CoV-2 genome stability for further *in-vivo* immunogenicity evaluation in HLA-A2.1 transgenic mice. Mice received a single subcutaneous injection of each peptide combined with the universal PADRE helper T-cell epitope^30^ and emulsified in Montanide ISA-51 adjuvant. Immunogenicity was assessed in the spleen and draining lymph nodes 15 days after vaccination by ex-vivo ELIspot restimulation with the corresponding WT peptide to evaluate cross-reactivity of elicited T cell response towards WT epitopes. IFNγ ELIspot after CD8 T cells restimulation showed large immunogenicity in-vivo responses elicited by single vaccination with 35 out of 60 (58%) positive CD8 T cells epitopes derived from all of the 11 selected proteins (**Figure 3A**). Similarly, broad immunogenicity response was observed by HLA-A*0201-tetramer flow cytometry analyses to a higher number of epitopes since 44 out of 60 (73%) evaluated peptides, derived from 10 out of the 11 selected proteins, exhibited significant frequency (0.1-1%) of viral-specific CD8 T cells (**Figure 3B**). Phenotypic characterization of Tetramer+ cells showed that 17 out of 44 (39%) positive peptides elicited viral-specific CD8 T cells with mainly a Trm phenotype: co-expressing the memory marker CD44, the CD103 αE integrin, the Th1-biased CXCR3 chemokine receptor and to a lesser extent the CD49a α1 integrin (**Figure 4**). Altogether, these data showed that optimized peptide vaccination against selected SARS-CoV-2 epitopes elicits robust and broad Th1-biased immunogenicity against several structural (S, M, N) and non-structural proteins in HLA-A2 expressing mice and that several peptides induce viral-specific memory CD8 T cells displaying all characteristics of T lymphocyte sentinels in barrier tissues.

**Figure 3:**
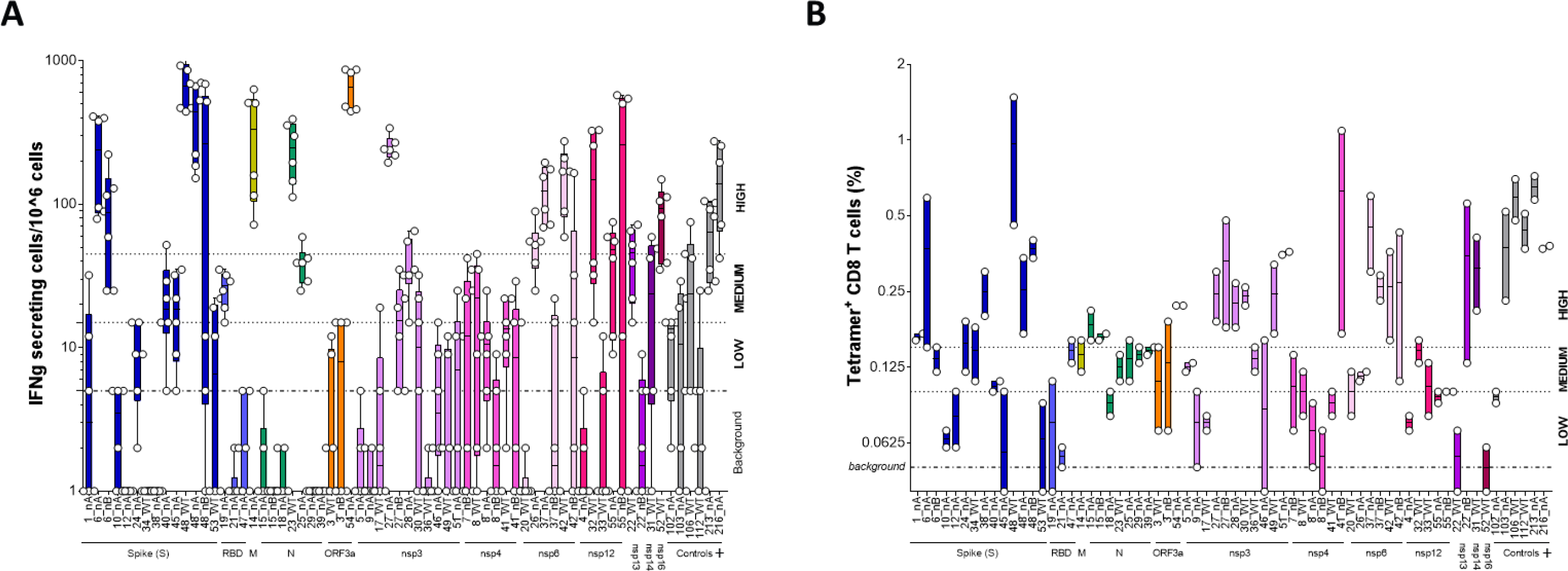
T-cell epitope vaccination elicits broad SARS-CoV-2 protein immunogenicity in-vivo. (**A**) IFNγ Elispot responses of pooled spleen and draining lymph nodes cells were restimulated for 24 hours with each of the isolated peptides. Data are mean +/- SEM (n=6). Medium level: Three-fold background. High level: Nine-fold background (**B**) Frequency of Tetramer^+^ CD8 T cells (gating strategy in Figure 4A). Medium and High response threshold were defined based on 2-fold or 3-fold increase respectively compared to the background frequency measured in the non-vaccinated mice control group. Controls+ are MEMOPI^®^ peptides with previously validated immunogenicity. Data are mean +/- SD of pooled female (n=3) and pooled male (n=3) vaccinated mice. Medium level: Two-fold background. High level: Three-fold background

**Figure 4:**
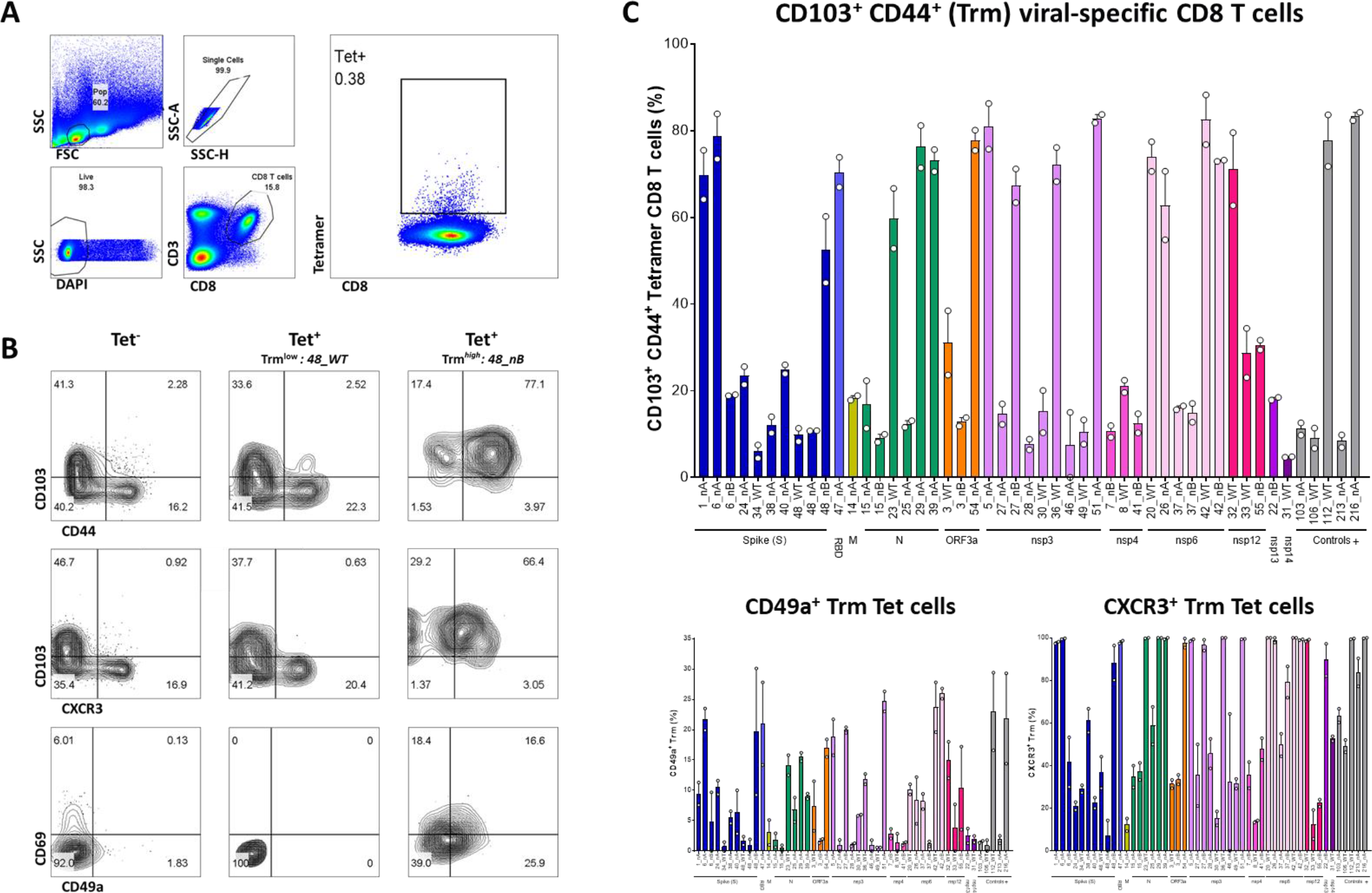
T-cell epitope vaccination elicits Tissue-resident memory viral-specific CD8 T cells. (**A**) Flow cytometry gating strategy to define Tetramer^+^ CD8 T cell responses. (**B**) Representative example and gating strategy of Trm phenotype based on CD44, CD103, CXCR3, CD49a and CD69 expression in Tetramer^+^ CD8 T cells as compared to Tetramer^-^ CD8 T cells. (**C**) *Top:* Frequency of CD103^+^ CD44^+^ (defined as Trm) cells within Tetramer^+^ CD8 T cells (defined as medium or high response based on Figure 3). *Bottom:* Frequency of CD49a^+^ or CXCR3^+^ cells within Trm^+^ Tetramer^+^ CD8 T cells. Controls+ are MEMOPI^®^ peptides with previously validated immunogenicity. Data are mean +/- SD of pooled female (n=3) and pooled male (n=3) vaccinated mice.

### CD8 T-cell epitopes ex-vivo reactivity in asymptomatic and convalescent COVID-19 individuals compared to unexposed healthy donors

In order to identify and select naturally SARS-CoV-2 CD8 T cell immunodominant epitopes, peripheral blood mononuclear cells (PBMC) from asymptomatic and moderate or severe COVID-19 patients with a previously confirmed (at least one month before sampling) and recovered SARS-CoV-2 infection were restimulated ex-vivo for one week with each of the 60 selected peptides derived from 11 proteins of SARS-COV-2. IFNγ responses over 48 hours of restimulation with HLA-A2+ Tap-deficient (T2) human cells were analyzed and compared to IFNγ responses of of unexposed healthy donors. The serology of all unexposed healthy donors was negative while all asymptomatic, moderate and severe COVID-19 individuals were IgG+ for anti-Spike antibodies. As recently reported for CD4+ T cells response^64,65^, we first observed some positive IFNγ response in unexposed donors to few CD8 T cells epitopes, in particular for those derived from structural Spike and N proteins (**Figure 5**). No response to the Spike_RBD or M structural proteins were observed in unexposed individuals, while CD8 T cell responses to non-structural proteins were limited to nsp12 and nsp13. In contrast, broad positive IFNγ response was reproducibly measured against 43 out of 60 selected CD8 T cells epitopes derived from all evaluated SARS-CoV-2 proteins (n=11) in convalescent subjects of moderate or severe COVID-19 as well as in asymptomatic individuals but to a lesser extent.

**Figure 5:**
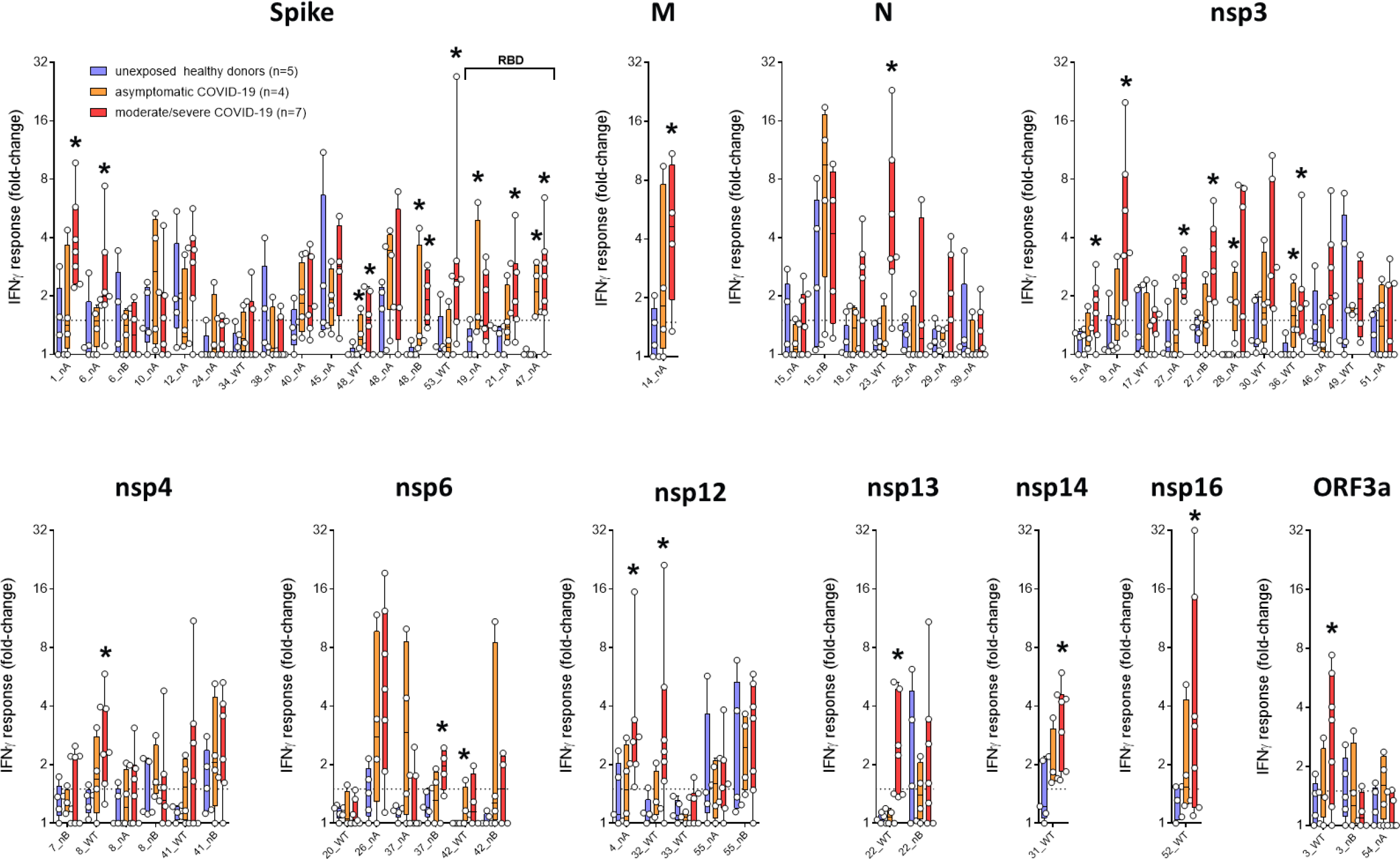
Ex-vivo T-cell epitope reactivity in asymptomatic and convalescent COVID-19 individuals and unexposed donors. IFNγ secretion responses for 48 hours after one week of restimulation of human PBMC from unexposed HLA-A2+ healthy donors (n=5), asymptomatic confirmed COVID-19 HLA-A2+ individuals (n=4) and moderate or severe COVID-19 HLA-A2+ convalescent patients (n=7) with each of the isolated peptides and HLA-A2+ antigen-presenting cells. Data were normalized to negative control peptides. Data are expressed as mean +/- min to max. *p<0.05

Altogether, when we compared convalescent SARS-CoV-2 individuals to unexposed healthy donors, we identified 25 significantly different CD8 T cell immunodominant epitopes against 3 structural proteins (S, M, N), 1 accessory factor (ORF3a) and 7 non-structural proteins (nsp3, nsp4, nsp6, nsp12, nsp13, nsp14, nsp16). 16 of these epitopes are of particular interest for vaccination since they were able to elicit also in-vivo immunogenicity (Elispot response) against all 11 structural and non-structural SARS-CoV-2 proteins after a single peptide injection. Finally, we selected a combination of 12 CD8 T cells epitopes based on manufacturing facilities, HLA-I coverage, previous CoVs homology and SARS-CoV-2 proteins diversity considerations (**Table 2**). These 12 epitopes covered the 11 selected proteins, 1 epitope/protein excepting Spike for which 2 epitopes (including 1 RBD epitope) have been selected. Bioinformatic analyses illustrate these 12 epitopes are not restricted only to the HLA-A*0201 allele, hence are predict (netMHCpan score < 1) to bind efficiently to different HLA-I (A, B, C) alleles with high genetic coverage in all geographical region of the world. Despite HLA polymorphism and different worldwide HLA-I distribution, the combination of these 12 T cell epitopes should induce at least 1 to 4 positive peptide responses in all individuals globally and achieve the 60-70% ‘herd immunity’ threshold^66^ with at least 4 to 5 positive peptide responses in each geographical region (**Table 3**).

**Table 2:**
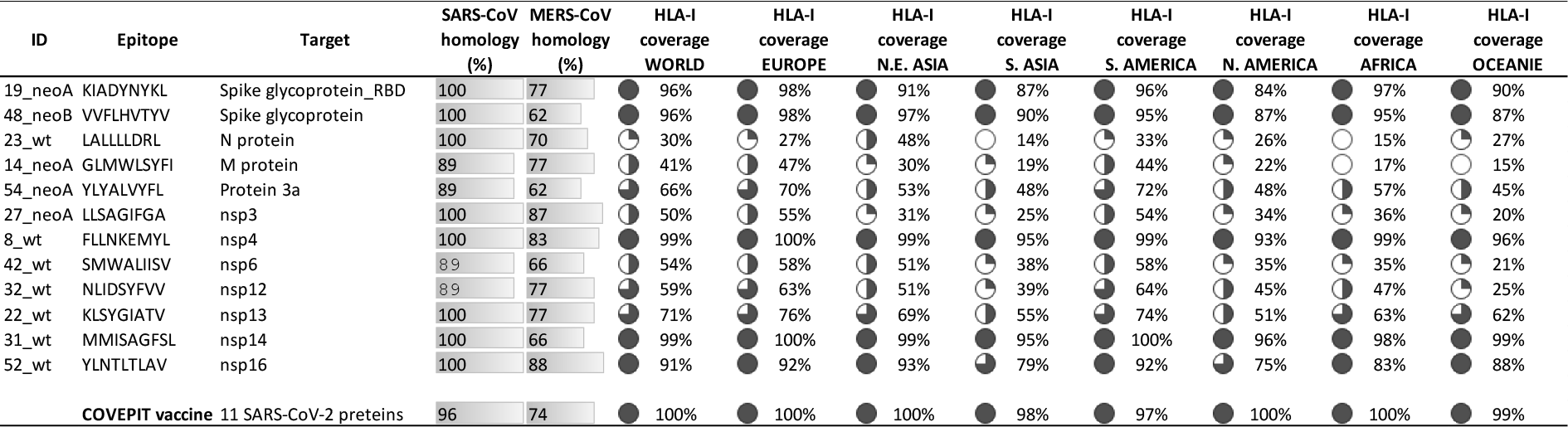
Selected CD8 T-cell epitopes, previous CoVs homology and HLA-I coverage per geographic region.

**Table 3:**
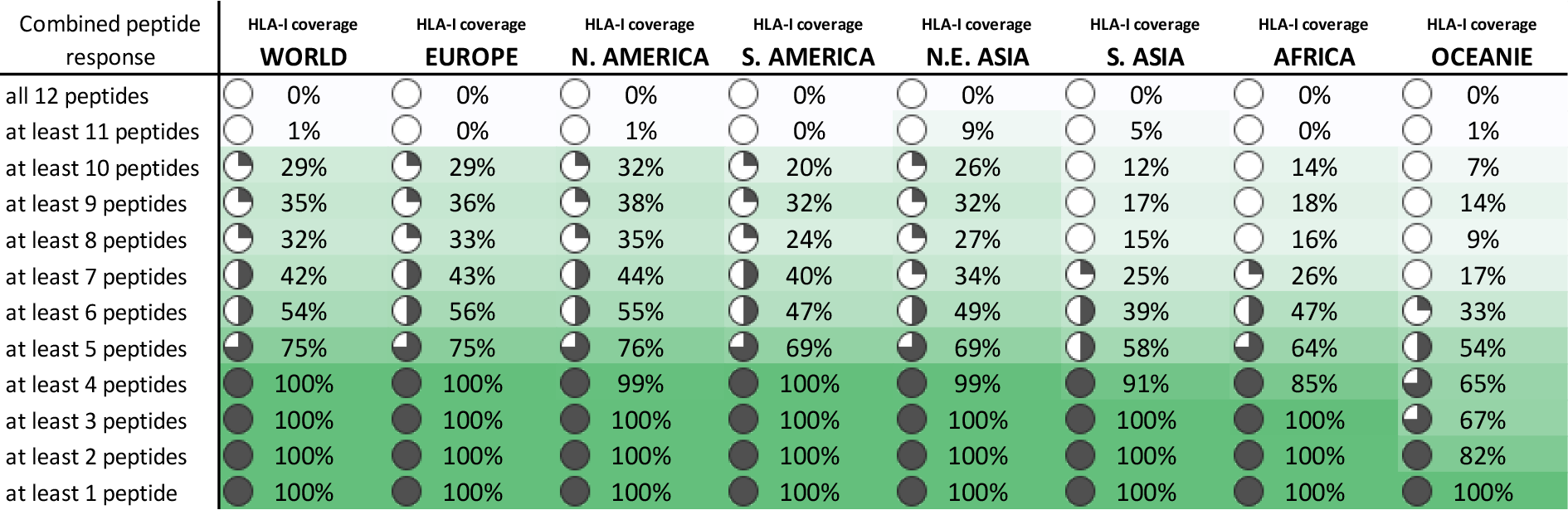
Responses to peptides combinations depending on HLA genomic frequency per region.

## Discussion

Here we report a differentiated SARS-COV-2 vaccine design based on memory T-cell induction technology. Using sequence design through reverse vaccinology selection approach based on previous CoVs knowledge on immunodominant epitopes and computational immunology optimization, we developed a combination of 12 CD8 T cell synthetic peptides originating from 11 SARS-CoV-2 structural and non-structural proteins capable to cover HLA polymorphism with high coverage globally and to induce immunogenicity to different proteins independently of HLA alleles expression. These epitopes are naturally immunogenic after SARS-COV-2 infection in recovered individuals and, for most of them, elicit a specialized sub-population of viral-specific memory CD8 T cells with a tissue-resident phenotype hence capable to migrate, stay attached and patrolling in airways barrier tissues.

Current vaccine approaches using various technology have already showed great anti-Spike antibody induction in Phase 1 and Phase 2 clinical trials as well as some interesting effector T cell responses against the vaccine antigen however without any indication of lasting memory T cell response ^67–70^. Furthermore, this first generation of vaccines are all monovalent, targeting only the Spike or a part of the Spike structural protein and built in urgency using the first Wuhan SARS-CoV-2 sequence. Genomic diversity and hotspot mutations analyses in up to 30000 SARS-CoV-2 genomes, illustrated that the virus had undergone mutations several times already in the last months^71^, adapting itself to its new host, and that new strains bearing significant Spike mutation (e.g. D614G) have emerged, spread globally and became the most prevalent form^63^. Once a mutated virus infects the host cells by escaping the antibodies, it then relies upon the T cell mediated immunity to fight against the virus. Viral proteins are processed into short peptides inside the infected cells and then loaded onto major histocompatibility complexes (MHC) proteins. After that, the MHC-peptide complexes are presented on the infected cell surface for recognition by specific T cells. For a universal vaccine, it is hence essential to understand genetic evolution and stability of the virus and to select immunodominant T cell epitopes that originate from conserved regions of the virus covering all circulating strains of SARS-CoV-2 and anticipating future mutation and evolution of the virus which appear more likely in hotspot regions. Similarly, multi-target vaccine are less susceptible to antigenic-drift and virus evasion after immune pression to a single antigen such as with monovalent vaccine.

The discovery of T cells as residents in tissues has initiated a paradigm shift in the way we study and understand T cell–mediated immunity: from a circulating and surveilling population transiting through barrier tissues to patrolling sentinels at the interface with the outside environment^15–17^. Trm act as immune sentinels and rapidly respond to infection by orchestrating local protective immune responses to eliminate pathogens. Because of their localization directly in tissues, they are able to immediately recognize infected cells or their cognate antigens presented by antigen-presenting cells. Trm protective immunity have been demonstrating to a number of infectious pathogens (viruses, bacteria, fungi, and parasites) in diverse sites such as mucosal and barrier sites (lung, skin, and intestines). In the lung, Trm mediate optimal protection against respiratory viruses such as influenza and respiratory syncytial virus (RSV), whereas circulating memory T cells and neutralizing antibodies appear to be dispensable^21,72–74^ Numerous findings indicate now the importance of both CD4+ and CD8+ Trm in tissue-localized immunity to pathogens, with CD8+ Trm more prevalent in viral and parasitic infections whereas CD4+ Trm mediate protection to bacterial and fungal invaders^17^. Furthermore, the CXCR3 chemokine receptor have been reported to be expressed on human lung T-cells and demonstrated to play a significant roles in T-cell homing to the lung parenchyma and ariway since CXCR3 inhibition prevents airway immune responses and vaccine-induced T cell entry to the lung mucosal compartments^75–79^. Here we showed that, as previously observed with our MEMOPI^®^-based neoepitope cancer vaccine approach, several peptides induced viral-specific CD8 T cells expressing the E (CD103) and α1 (CD49a) integrins and the CD44 memory marker, altogether characteristic of Trm. These cells express also the CXCR3 marker of chemoattraction, which is also a surrogate marker of Th1 CD8 T cells since CXCR3 is transactivate directly by the Th1 master gene T-bet^79^. CD103 and CD49a expression are key molecules in Trm biology since providing them the capacity of interaction, adherence and mobility with mucosal tissues, in particular epithelial cells and collagen directly underlying epithelial cells^80,81^.

To summarize, this differentiated Th1-biased memory CD8 T cell SARS-CoV-2 vaccine has been developed using reverse vaccinology selection approach, computational immunology optimization and synthetic peptide synthesis technology. The development of inactivated virus vaccine, besides Good Manufacturing Practices (GMP) system, require extremely high manufacture standard to avoid medical accidents due to the failure of complete inactivation of virus toxicity. Furthermore, different from nucleic acid (e.g. RNA) vaccines, peptide vaccine has relatively more mature manufacturing process. The successful launch of previous peptide vaccines has already demonstrated the safety and efficacy of peptide-based vaccines. The first generation of SARS-CoV-2 vaccine has been based on the first SARS-CoV-2 Wuhan sequence and target only one antigen. Here we propose a second generation of SARS-CoV-2 vaccine taking into consideration virus genetic evolution these last 6 months after global spreading and which targets different structural and key non-structural proteins in stable genome sequence region to cover all circulating SARS-CoV-2 strains and anticipate future virus evolution. Selected T cell epitopes display also very high homology with previous pandemic coronavirus (SARS-CoV-1 and MERS-CoV) and hence may be suitable and ready for future anticipated coronavirus pandemic infection. These preclinical findings on Tissue-resident memory CD8 T-cell responses, providing long-term immunity, were observed after a single injection of a multi-target COVID-19 vaccine. The validation in human PBMC from convalescent asymptomatic, moderate and severe COVID-19 patients, support further development of this peptide vaccine anticipating future evolution of the virus in hotspots mutation regions.

## Methods

### Study Approval

Animal housing and procedures have been conducted according to the guidelines of the French Agriculture Ministry and were approved by the regional ethical committee (APAFIS 25256.) as well as according to the guidelines of Jackson Laboratory (Bar Harbor, USA) and approved by the Institutional Animal Care and Use Committee (IACUC #20031). Human studies were performed under the clinical protocol COVEPIT-1 approved by French Central Ethic Committee (CPP) and registered by the French Regulatory Authority (ANSM) under the ID-RCB n° 2020-A01654-35. Written informed consent has been obtained from each of the participating subject.

### HLA-A*0201-peptide binding evaluation

T-cell WT and mutated peptides binding property on HLA-A2 has been evaluated using the Flex-T HLA-A*02:01 monomer ultraviolet (UV) exchange assay according to the manufacturer recommendation (Biolegend, San Diego, USA). HLA-A*02:01 monomer (200 μg/ml) were exposed to a 366-nm UV lamp in the presence or absence of 400 μM of peptide. After UV-exposure, HLA-peptide complexes were incubated at 37°C for 30 min to promote unfolding of peptide-free HLA molecule. HLA-peptide complexes stability was detected by ELISA with β2-microglobulin coated antibodies and incubation of 3 ng/ml of complexes for 1h at room temperature under shaking condition. Avidin-HRP were used to reveal stable biotinylated HLA-peptide complexes and absorbance was monitored at 450 nm. Data are expressed as percentage of binding relative to an MEMOPI^®^ internal positive control neoepitope.

### SARS-CoV-2 genome diversity evaluation

A visualization tool was used to determine T-cell and B-cell epitope location in SARS-CoV-2 genomes according to single nucleotide polymorphism (SNPs) and homoplasic site (*https://macman123.shinyapps.io/ugi-scov2-alignment-screen/*)^85^. 46723 SARS-CoV-2 genomes isolated from patients worldwide were aligned against the Wuhan-Hu-1 reference genome NC_045512.2. A total of 12706 SNPs has been identified corresponding to 398 homoplasic sites with recurrent mutations. Peptides have been blasted with tblastn algorithm against the Wuhan-Hu-1 reference genome NC_045512.2 to determine the nucleotide coordinates for each peptide. The online tool was then used to identify the peptides corresponding to a homoplasic site.

### T-cell epitopes immunization

B6.Cg-*Immp2l_Tg(HLA-A/H2-D)2Enge_*/J (HLA-A2.1) transgenic mice (JAX # 004191, The Jackson Laboratory, Bar Harbor, USA) received a single subcutaneous injection of 6 SARS-CoV-2 peptides (50μg each, WT and mutated peptide of a same epitope have not been evaluated in same mice) plus the universal PADRE helper T-cell epitope^30^ emulsified in Montanide ISA-51 adjuvant. Immunization was measured 15 days after injection. 3 males and 3 females have been evaluated per group. Freshly harvested spleen and draining lymph nodes have been pooled by sex per group and analyzed by flow cytometry analyses. CD8+ T cells have been isolated using MACS microbeads and restimulated individualy with each evaluated peptide. The frequency of IFNγ-secreting CD8+ T cells was measured by ELIspot in parallel of tetramer staining for each peptide evaluated by flow cytometry.

### COVID-19 patient cohorts

All subjects were enrolled in the COVEPIT-1 clinical trial, an open-label, multicentric and prospective study with minimal risk and constraints designed to assess the memory T cell in subjects who recovered from COVID-19. Subjects were enrolled at the Groupement Santé du Bataillon des Marins-Pompiers (Marseille-France) and GHR MSA– Hôpital Emile Muller (Mulhouse, France). The main objective was to test the subject’ memory T cells reactivity to a selection of SARS-CoV-2 antigens. Enrolled subjects must have a proven COVID-19 infection which recovered 1 to 6 months before study entry. Eligible subjects are male and female of 18 to 70 years old diagnosed for COVID-19 using a PCR test from a nasal and/or oropharyngeal swab, and/or a serological test, and/or a chest CT with lesions suggestive of COVID-19. Subjects were excluded if pregnant or breastfeeding, unable to fulfill the protocol requirement, with an history of cancer 5 years prior to study entry (except for localized or in situ cancer), history of head injury or sepsis 1 year prior to study entry, chronic infections (e.g. HIV infection, chronic hepatitis B, active viral hepatitis C or bacterial or fungal infection) requiring a systemic treatment in the month prior to COVID-19, disease (auto immune or inflammatory disease, transplant recipients…) requiring a immunosuppressive or immunomodulator treatment and/or, corticosteroids at an equivalent dose of prednisone >10mg/d for more than 15 days or > 40 mg/d for the last 15 days prior to COVID-19; corticosteroid during COVID-19 were not considered as an exclusion criterion.

### Ex-vivo PBMC restimulation with SARS-CoV-2 peptide

PBMC were isolated after a Ficoll density-gradient centrifugation and a red blood cell lysis. HLA-A2 phenotyping was performed by flow cytometry (clone BB7.2, BD Bioscience). Ex-vivo stimulation protocol was adapted from a previously described protocol^86^. HLA-A2+ positive PBMC (10^6^/well) were incubated in RPMI 1640 containing 10 mM HEPES, 2 mM L-glutamine, 1mM Sodium Pyruvate, 2% human AB serum, 10% bovine serum and non-essential amino acids in 48-well plates. During the first week of culture, PBMC were cultured with 3μg/mL of each isolated peptides and IL-21 (30ng/mL; Miltenyi, Paris France). Fresh medium containing IL-21 (30 ng/mL), IL-7 (5 ng/mL; BioRad, Paris France), and IL-2 (10 ng/mL; Miltenyi, Paris France) and peptide-loaded HLA-A2+ Tap-deficient (T2) cells were added to the culture for the next two days. Ex vivo T-cell viral stimulation was evaluated by IFNγ supernatant quantification (BD Biosciences, US). The percentage of background IFNγ secretion was determined by the response of PBMC co-cultured with non-loaded T2 cells and negative control peptide, then fold change was calculated over the IFNγ secretion background for each donors.

### Statistical analysis

Continuous variables were expressed as the mean ± SEM, unless otherwise indicated, and raw data were compared with nonparametric tests: Mann-Whitney for 2 groups or Kruskall-Wallis with Dunn’s comparison when the number of groups was > 2. P values of <0.05 were considered statistically significant. All statistical analyses were performed on GraphPad Software (GraphPad Software, San Diego, CA).

## Acknowledgments

This work was supported by funding from Nantes Metropole as part of the Metropolitan Fund to Support Health Innovations Linked to the COVID-19 Health Crisis. We thank clinicians and patients involved in the COVEPIT-1 trial as well as the eXYSTAT company for biometric expertise and data management for the COVEPIT-1 trial.

